# Studying the evolutionary potential of ancestral aryl sulfatases in the alkaline phosphatase family with droplet microfluidics

**DOI:** 10.1101/2024.12.10.627700

**Authors:** Bernard D. G. Eenink, Josephin M. Holstein, Magdalena Heberlein, Carina Dilkaute, Joachim Jose, Florian Hollfelder, Bert van Loo, Erich Bornberg-Bauer, Tomasz S. Kaminski, Andreas Lange

## Abstract

Characterizing the dynamics and functional shifts during protein evolution is essential, both for understanding protein evolution and for rationalizing efficient strategies for e.g. enzymes with desired and effective functions. Most proteins organize in families, sets of divergent sequences which share a common ancestor and have a similar structural fold. We study aryl sulfatases (ASs), a subfamily of the large and evolutionary old alkaline phosphatase superfamily. In this manuscript we present how ultrahigh-throughput droplet micro-fluidics can be used for studying aryl sulfatases and their computationally reconstructed putative common ancestors. We compare the evolvability and robustness of three ancestors and four extant ASs which all exhibit catalytic promiscuity towards a range of substrate classes. Fourteen libraries with varying mutation rates were expressed in single-cell microdroplets. In general, higher mutation rates resulted in wider distribution of active variants but fewer improved variants overall. However, the impact of mutation rate differed between enzymes, with some benefiting from higher and others from lower mutation rate, underscoring the need to test diverse mutagenesis regimes.

## Introduction

A fundamental aspect of understanding the evolution as well as the structural and functional connections of proteins involves categorizing proteins into families [1]. Essentially, protein families are believed to have originated from a shared ancestor through gene duplication, followed by sequence and functional divergence [2, 3], with structural conservation generally being more pronounced [4, 5]. Conversely, to some extent, this apparent conservation arises from structural limitations, which constrain the extent at which sequence modifications can occur before a protein loses its functionality.

Enzyme promiscuity is now widely recognized [6, 7] as a key factor in the evolution of new functions, providing organisms with moderate activity toward non-standard substrates and lowering the threshold for selective advantage [7, 8, 9]. Promiscuous enzymes serve as promising starting points for evolving new activities, potentially even before gene duplication [7], enabling the rapid development of biocatalysts for green chemistry and xenobiotic degradation [10].

Directed evolution enhances protein functions through iterative mutation and selection, mimicking natural evolution [11, 12]. It is used to improve traits such as catalytic efficiency [13], stability [14], enantioselectivity [15], and tolerance to alternative conditions [16], expanding the enzyme repertoire for novel applications. However, a successful campaign requires not only selecting a candidate based on kinetic parameters [10, 17] but also understanding the local fitness landscape, an essential yet more complex task. For example, shallow fitness peaks may be easier to reach and variants may be easier maintained due of the sheer number of viable variants in the near vicinity while steeper fitness peaks may be difficult to reach and variants more easily lost ("survival of the flattest") [18, 19].

### Aryl sulfatases

Aryl sulfatases (AS) are a family of the alkaline phosphatase (AP) superfamily. ASs catalyize the de-esterification of sulfate mono-esters [20]. In addition to this main reaction, many ASs show catalytic promiscuity towards other substrate classes [21]. Substrate classes can reach high levels, sometimes on par with the catalytic efficiency of specific enzymes towards a large number of substrate classes [22, 23], making ASs suitable for the investigation of the characteristics and evolvability of ancestral enzymes.

Microfluidics is capable of screening the 10^5^-10^6^ variant libraries [24] needed in a comprehensive manner. A limitation of microfluidics is that cell lysis is needed for the enzyme in the cell and the substrate in the medium to meet. Thus cells cannot be recovered after sorting, and the DNA of selected variants needs to be re-transformed [13].

A solution out of this dilemma has been recently proposed because active multimeric APs can be displayed on the cell surface of *E. coli* [25, 21], the multimeric AP model systems can be screened and recovered efficiently using ultra-high throughput library sorting methods [21]. Additionally, Model substrates for ASs are widely available, and can be used to detect sulfate, phosphate and phosphonate esterase activity using a variety of detection methods (Figure 1) [21].

**Figure 1:**
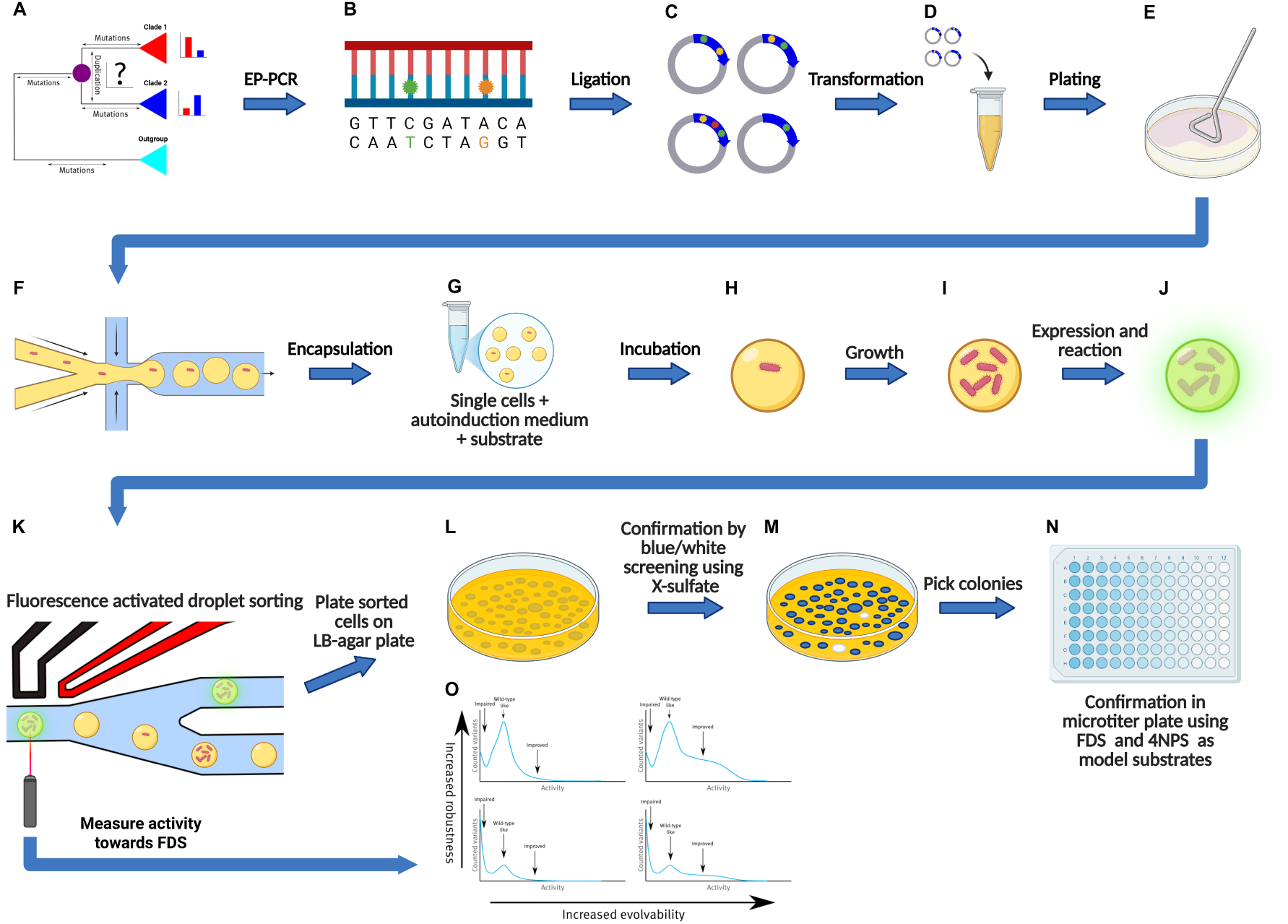
Overview of the protocol used. Ancestral enzymes were reconstructed using ASR (A). epPCR was used to generate mutant libraries of each enzyme (B). Variant libraries were ligated into pBAD-AT-shift in frame with the AIDA autodisplay construct (C). *E. coli* cells were tranformed with the libraries (D) and spread on agar plate. Grown colonies were subsequently scraped from agar plate (E) and encapsulated inside a water-in-oil microdroplet together with LB-medium and substrate (F). The water-in-oil droplets were collected in an eppendorf tube (G) and incubated (H) after growth and expression (I and J) droplets were sorted using fluorescence activated droplet sorting (K). The activity towards FDS in each droplet was measured (O). 0.01% of the most active variants were sorted, plated on LB agar plate (K) and confirmed using Blue-White screening with X-sulphate (**2a**) on agar plate and 4NPS (**1a**) and FDS (**3a**) in microtiter plate (N). This figure was prepared with BioRender.com.

### Ancestral sequence reconstruction

Ancestral sequence reconstruction (ASR) is used to deduce ancestral sequences at divergent nodes in a phylogenetic tree [26, 27]. Enzymes inferred through ancestral reconstruction tend to exhibit several noteworthy properties, such as high thermostability [28, 29, 30], catalytic promiscuity [29, 31], and retaining much higher activity at lower temperatures [32]. ASR studies into glucosidases [3] and malate- and lactate dehydrogenases [33] have shown minor molecular changes can drastically alter enzyme function. Increases in thermostability by 20-30 °C can be achieved [34, 35, 36]. However, whether this increase in thermostability reflects a genuine evolutionary trend or, to some extent, an artifact arising from the chosen reconstruction method remains to be seen [30, 37].

High thermostability and catalytic promiscuity make proteins ideal for directed evolution, as promiscuous enzymes adapt more easily to new functions, and thermostability allows function-enhancing but destabilizing mutations without compromising their function too much [38]. ASR expands the pool of templates beyond natural limits, enabling exploration of alternative evolutionary paths under different selection pressures [39]. Since ancestral enzymes can be readily expressed, investigating their sequence space alongside extant enzymes may reveal untapped catalytic potential. Caution should be taken when addressing specificity, as it can be seen on two levels; First at a Biological level, the affinity of a protein for potential targets at varying concentrations, as a result of selection. And second at the intrinsic level, a general affinity to targets not encountered and thus not the result of selection, that can nonetheless be a basis for novel functionality [40]. Studies in ancestral sequence reconstruction generally probe biological rather than intrinsic specificity, and biological specificity does not necessarily imply intrinsic specificity [41, 40].

### Micro-droplet sorting

To screen large amounts of enzyme variants, high-throughput screening methods are required. Water-in-oil emulsion-based micro-droplet sorting enables the screening of large libraries of enzymes and other proteins. The reaction volume is reduced to picoliters, increasing throughput and minimizing reagent usage [24, 42, 43]. By ensuring single occupancy each microdroplet functions as a mini-bioreactor, hosting one cell, or even one plasmid, that produces a single enzyme variant, catalyzing substrate turnover, thus maintaining genotype-phenotype linkage. Droplet sorting has been widely used to screen various classes of enzymes with ultra-high throughput, with both in vivo and in vitro expression systems [44, 45]. The most straightforward way to mix enzymes and substrate is cell lysis, after which the expression vector is recovered and sequenced or re-transformed into the host organism [43, 24]. To directly recover cells after sorting enzyme can be autodisplayed on the cell surface and lysis omitted [25, 46, 13].

Recent studies show cell growth inside droplets can improve sensitivity and recovery while preserving genotype-phenotype linkage [47, 48]. In these methods, microdroplets with growth medium are inoculated with single cells, which multiply to form monoclonal populations. Cell surface display relocates the enzyme from inside the cell to the surface, where it can interact with substrate in the medium, omitting a lysis step and allowing for direct recovery by plating sorted cells on agar plates.

In this study we use the *E. coli* autodisplay system (Figure 2) [46, 25], which presents enzymes on the outer membrane. This enables screening and sorting of enzyme libraries with intact, viable cells in microdroplets. Previously, this technique improved the catalytic efficiency of *Sp*AS1 by up to 6.2-fold towards 4-nitrophenyl sulfate and up to 30-fold towards fluorescein disulfate in a single round of directed evolution [13]. The autodisplay system has also demonstrated the ability to express properly folded, active dimeric enzymes [49, 50], including ASs [13].

**Figure 2:**
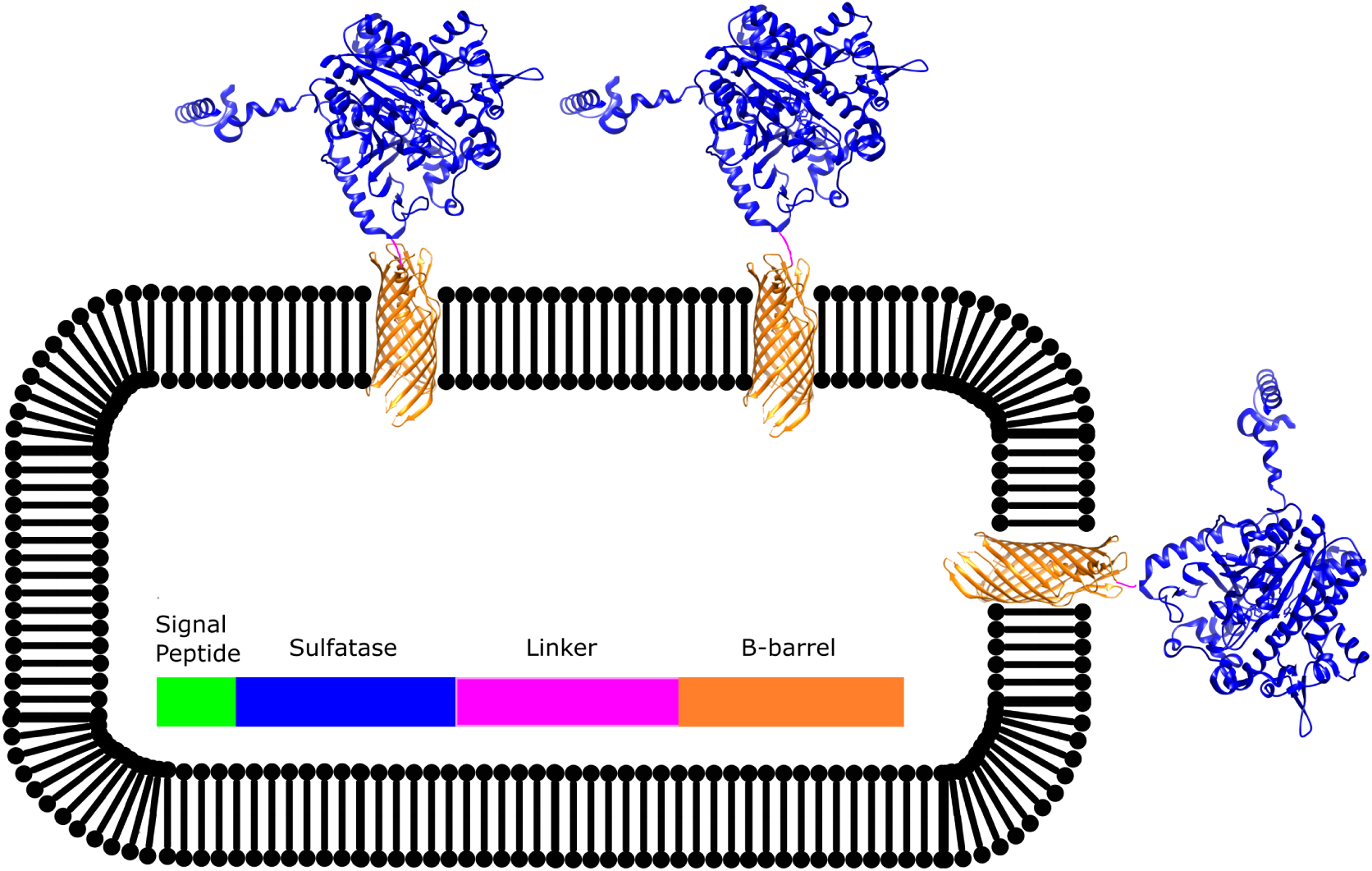
Schematic overview of the sulfatase enzymes displayed on the outer membrane of *E. coli* cells. A signal peptide directs the fused protein towards the periplasmic space where the AIDA β-barrel folds into the outer membrane and displays the passenger enzyme at the surface [48, 46].

### Aims

This study aims to directly compare ancestral and modern AS enzymes from the AP superfamily using a unified experimental approach. By integrating droplet microfluidics, the *E. coli* autodisplay system [25, 46], and ancestral sequence reconstruction (ASR), we analyze aryl sulfatases and phosphonate monoester hydrolases from the alkaline phosphatase superfamily [21]. This framework enables broader enzyme characterization, with the following objectives: (i) systematically assessing the evolvability (proportion of variants with improved catalytic efficiency over the wild type) and robustness (proportion of variants retaining wildtype-like or better catalytic efficiency) of ancestral proteins, (ii) correlating catalytic efficiency gains observed in high-throughput screens with those in cytosolically expressed, purified proteins, and (iii) comparing the local fitness landscapes of extant and ancestral AS enzymes.

## Materials and Methods

### Ancestral reconstruction procedure

The multiple sequence alignment calculation as well as the phylogeny reconstruction were performed based on the amino acid sequences. For the ancestral gene reconstruction, the coding DNA sequences of each protein in the dataset were downloaded from the NCBI RefSeq database [51] and aligned similarly to the previously calculated protein sequence alignment using TranslatorX server [52]. The resulting nucleotide sequence alignment and phylogenetic tree were used as input for the codeml program of the Phylogenetic Analysis by Maximum Likelihood (PAML) software package [53]. The ancestral nucleotide sequences were reconstructed based on coding DNA sequence alignment. The manual refinement of ancient sequences was based on the maximum parsimony principle, assuming the lowest possible number of changes. Each position of the final alignment was manually inspected in terms of the amino acids presence/absence in the reconstructed ancestor versus its descendants. The sequence similarity between the reconstructed and the extant proteins was calculated using blastp, using the presented dataset as a local database. For subsequent cloning experiments, all *Xho*I, *Kpn*I, *Pst*I, *Bam*HI and *Hind*III restriction sites were removed from the reconstructed sequences. The sites were changed by substituting the third base of the codon encompassed in the restriction site to the second most probable base as reconstructed by PAML. At the C-terminus of all sequences, a TAA stop codon was added as the one with highest frequency in *E. coli*-61% cases according to the GenScript CodonUsageFrequency Table Tool available at http://www.genscript.com/cgibin/tools/codon freq table (accessed 2014/9/15) / 64% cases according to the Codon Usage Database [54]. To facilitate cloning, linker sequences were added at the N- and C- termini of each reconstructed gene (N-terminus: 5-GTACCCGGGGATCCCTCGAG-3,C-terminus: 5-CTGCAGGGGGACCATGGTCT-3). The sequences were optimized according to the Gen9 synthesis guideline. The BsaI and AarI recognition sites were removed (in the same way as other restriction sites) and the local GC content was adjusted using the Gen9 online optimization tool. The synthetic genes were purchased from Gen9.

### Cloning procedure for cytosol expression

For the cytosolic expression and purification of enzymes, DNA sequences were cloned into the pASKiba5+ vector (Figure 3). Ancestral sequences (Anc497, Anc498, Anc499) were cloned using the restriction sites *Pst*I+*Xho*I, while extant sequences (*Sp*AS2, *Sa*AS, *Ak*AS, *Rp*AS) were cloned using the *Xho*I+*Kpn*I restriction sites. Subsequently, chemically competent *E. coli* TOP10 cells (INOUE method) were transformed with the assembled plasmid.

**Figure 3:**
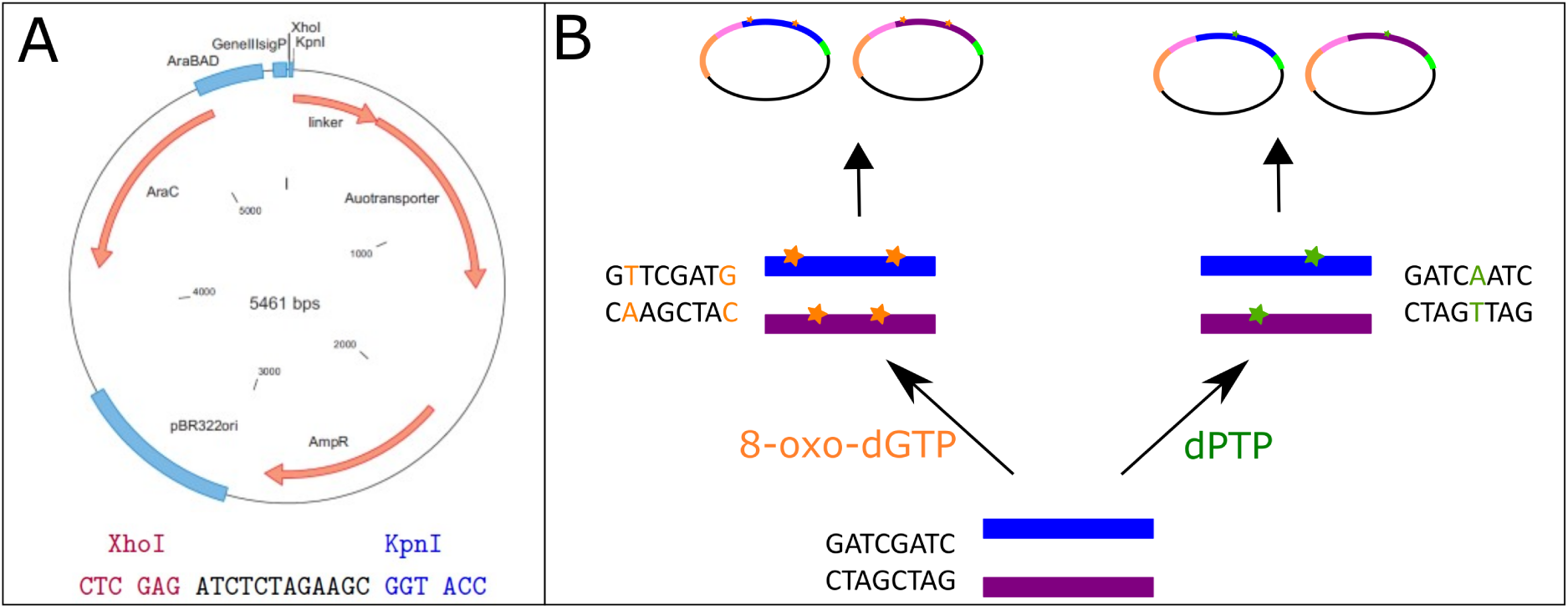
Overview of plasmid engineering used in this study. pBAD-AT-shift (A) was used for all vector engineering. epPCR libraries were generated by spiking the PCR reaction with the synthetic nucleotides 8-oxo-dGTP (yellow) and dPTP (green), which each have a unique substitution profile and a different mutation rate (B).

### Purification of cytosolically expressed enzymes

10 mL of LB-medium containing 50 µL/L ampicilin was inoculated with *E. coli* cells containing an expression vector for the respective ancestral and extant genes. This pre-culture was grown overnight at 37 °C. 2 L 2x YT medium (16 g/L tryptone, 10 g/L yeast extract, 5 g/L NaCl) was inoculated with the preculture and incubated at 37 °C while shaking until an OD_600_ of between 0.6 and 0.8 was reached. Expression was induced by adding anhydrotetracycline (200 µg/L followed by overnight growth at 25 °C. Cells were harvested by centrifugation, resuspended in HCl buffer pH 8.0 with EDTA-free Protease Inhibitor Cocktail (Roche). Cells were subsequently lysed by addition of 1x Bugbuster containing Lysonate and incubated for 1h at room temperature. Cell debris was removed by centrifugation (90 min, 15000x g, 4 °C), the supernatant was passed through a 0.45 µm syringe-driven filter. The proteins were purified using afinity chromotography with a Strep-Tactin column (IBA). Peak fractions were combined, concentrated down to 100 µM. The buffer was exchanged to 50 mM Tris-HCl pH 8.0 by passing the protein solution through PD MiniTrap G-25 columns (GE Healthcare). Protein concentration was determined from absorption at 280 nm. The extinction coe cients and molecular weights (MW) were calculated using ProtParam at ExPASy (http://web.expasy.org/protparam/). Protein aliquots were either used immediately or frozen in liquid nitrogen and stored at -80 °C.

### Reagents

4-NPS substrate was from Thermo-Fisher. FDS substrate was synthesized as previously described [55]. X-sulfate substrate was from Goldbio.

### Cloning procedure for autodisplay

For the expression of individual variants using autodisplay, DNA sequences were cloned into pBAD-AT-shift vector [56] (Figure 3) using the XhoI+KpnI restriction sites. Subsequently, chemically competent (INOUE method) *E. coli* E. cloni 10G cells were transformed with the assembled plasmids.

### Library generation methods

Libraries were created using error-prone PCR. In both reactions a mutagenic nucleotide analog, 2’-deoxy-P-nucleoside-5’-triphosphate (dPTP) or 8-oxo-2’-deoxyguanosine-5’-triphosphate (8-oxo-dGTP), respectively, was supplemented into a standard Taq DNA polymerase reaction (GoTaq, Promega). These nucleotide analogs can form pairbonds with multiple nucleotides and can be erroneously incorperated during PCR, when these nucleotide analogs get duplicated again, a different nucleotide may be incorperated in their place. Of these synthetic nucleotides, dPTP causes mainly transitions, while 8-oxo-dGTP causes mainly translations [57]. Subsequently a recovery PCR, using the same protocol but not incorperating synthetic nucleotides, was performed on the obtained PCR fragments to remove the incorporated nucleotide analogs. Sequences were then cloned into pBAD-AT-shift using the *Xho*I+*Kpn*I restriction sites (Figure 3). The average mutation rates were verified using Sanger sequencing as in [13].

### Expression of autodisplay constructs

Either glycerol stocks or single colonies on agar plate were used to inoculate 3-10 mL LB medium (100 µg/mL ampicillin) and incubated overnight at 37°C. Overnight cultures were used to inoculate fresh LB medium and incubated at 37°C until an OD_600_ of +-0.5. After reaching OD_600_ cells were transferred to expression temperature and incubated for another hour. Cells were then induced by addition of 0.02% L-arabinose and expressed overnight. After overnight expression cells were harvested by centifugation (5 min, 4000*g), resuspended in 1/3 volume SID buffer (pH 7.2, 7.6 or 8.0), according to previously determined pH optima [20, 58]. Cells were then washed three times by centifugation (5 min, 4000*g) and resuspended in fresh 1x SID buffer before measurements.

### Whole cell assays with autodisplayed variants

50-100 µl 1x SID buffer was added to 20-70 µL of resuspended cells for a total volume of 120 µL in a 200 µL microtitre plate. The cell density of each of the cells in each well was measured at OD_600_ before the addition of substrate. 80 µL of 2.5x final concentration **1a** (4NP) or **1b** (FDS) in 1x SID buffer was added for 200 µL final volume. The reaction is measured by increase of absorption value at (400 nm) **1a** (4NP) or fluorescence (λ_ex_ = 490 nm, λ_em_ = 515 nm).

### Microfluidic droplet screening

Frozen cells were resuspended in autoinduction medium (LB containing ampicillin 100 mg L^-1^), L-arabinose (0.04% (v/v) and 0.2% (v/v) D-Glucose. Cell concentration was adjusted to 0.3 cell per droplet volume based on the assumption that OD_600_ = 1 to correspond to 5×10^8^ *E. coli* cells mL^-1^. Droplets were produced in a double flow-focusing junction chip (design: https://openwetware.org/wiki/DropBase:droplet_generation_2_inlets, design iv: 40 µL). First, the autoinduction medium was mixed 1:1 with a solution of 40 µM fluorescein disulfate **1a** in SID buffer pH 7.5 just upstream the flow-focusing (FF) droplet generation junction. At the FF junction, fluorinated oil HFE-7500 (3M) containing 1% (v/v) fluorosurfactant-008 (RAN Biotechnologies) was used to break the continuous stream aqueous phase and generate monodisperse 40 pL droplets with an expected cell occupancy of λ = 0.35 (assuming Poisson distribution). Flow rates were following: 40 µL min^-1^ for both aqueous phases and 160 µL min^-1^ for the oil/surfactant phase. The droplet formation was monitored on an inverted microscope (SP981, Brunell Microscopes) equipped with a high-speed camera (Miro ex4, Phantom Research). The droplets were collected in a droplet chamber (as described in [59]). Generated droplets were incubated for 1 to 3 days at 30° C. During incubation the cells were aerated [47] by flushing with fluorinated oil HFE-7500 (3M) containing 1% (v/v) fluorosurfactant-008 at a rate of 4 µL per minute. After 2-3 days the droplets were sorted using FADS (as previously described in van Loo et al., 2019 [13], the LABVIEW script used is available online https://github.com/droplet-lab/spinDrop/tree/main/LabVIEW%20FADS. Droplets were re-injected from the droplet chamber onto a microfluidic chip (design: https://openwetware.org/wiki/DropBase:droplet_electrosorting_3) using HFE-7500 (3M) containing 1.5% (v/v) fluorosurfactant-008 (RAN Biotechnologies). Droplets were spaced with oil and pushed through a narrow detection channel to allow single droplet measurement. As the droplets were pushed past a 488 nm laser in the sorting Y-junction, the fluorescent activity inside each droplet as measured by recording emission at 497-553 nm to quantify substrate turnover. Droplets would then flow into a waste channel. When fluorescent activity exceeded a threshold (dependent on wild-type enzyme) pulse and function generators were triggered and generated a square pulse at 8 volts, which was amplified 100-fold by a high-voltage amplifier (610E, Trek) and applied on the sorting device via salt-water electrodes (5 M NaCl). This pulse pulled the selected droplets into a collection channel. The threshold was set to 0.1% of total droplets during a calibration and stabilization phase. Droplets sorted during this phase were discarded, once droplet flow was stabilized and threshold established, collection tubes were added to the collection channel and waste channel and droplet collection started. droplets were collected at a high frequency until a total of 1.000.000 droplets were sorted.

The droplets in the positive collection channel were collected in a solution of 100 µL Lucigen recovery medium, 45 µl HFE-7500 (3M) and 5 µL PFO surfactant. The collection tubing was flushed with HFE-7500 (3M) to collect all cells. An additional 500 µL of recovery medium was added after collection and the solution was mixed. The liquid phase was plated on LB+amp plates and incubated overnight.

### Blue-White screening

Cells, recovered from positive droplets after screening, were plated on agar plates (LB+amp) and incubated overnight at 37°C. Subsequently, cells were scraped from the plate and replated on a fresh LB+amp plate on top of a nitrocellulose filter. After overnight incubation at 37°C the nitrocellulose filter was transferred to a fresh LB+amp+arabinose plate and incubated overnight at room temperature to induce expression. The next day, the nitrocellulose filters were transferred to a fresh LB+amp+arabinose+X-Sulfate plate. Colonies capable of turning over the substrate shifter color to blue. Blue colonies bearing sulfatase activity were counted.

## Results and discussion

### Selection and confirmation of wild-type and ancestral aryl sulfatases

We reconstructed the aryl sulfatase (AS) and phosphonate monoesterase (PMH) sets of the alkaline phosphatase superfamily down to their last common ancestor ASPMH (Figure 4). We expressed several AS ancestors and characterized these ancestors in the lab. We decided to focus on the AS family as ancestral ASs were more amenable to hetererologous expression in *E. coli*, as well as a broader availability of model substrates. Thermostability of each wild-type was determined using a thermo-shift assay (Table S2)

**Figure 4:**
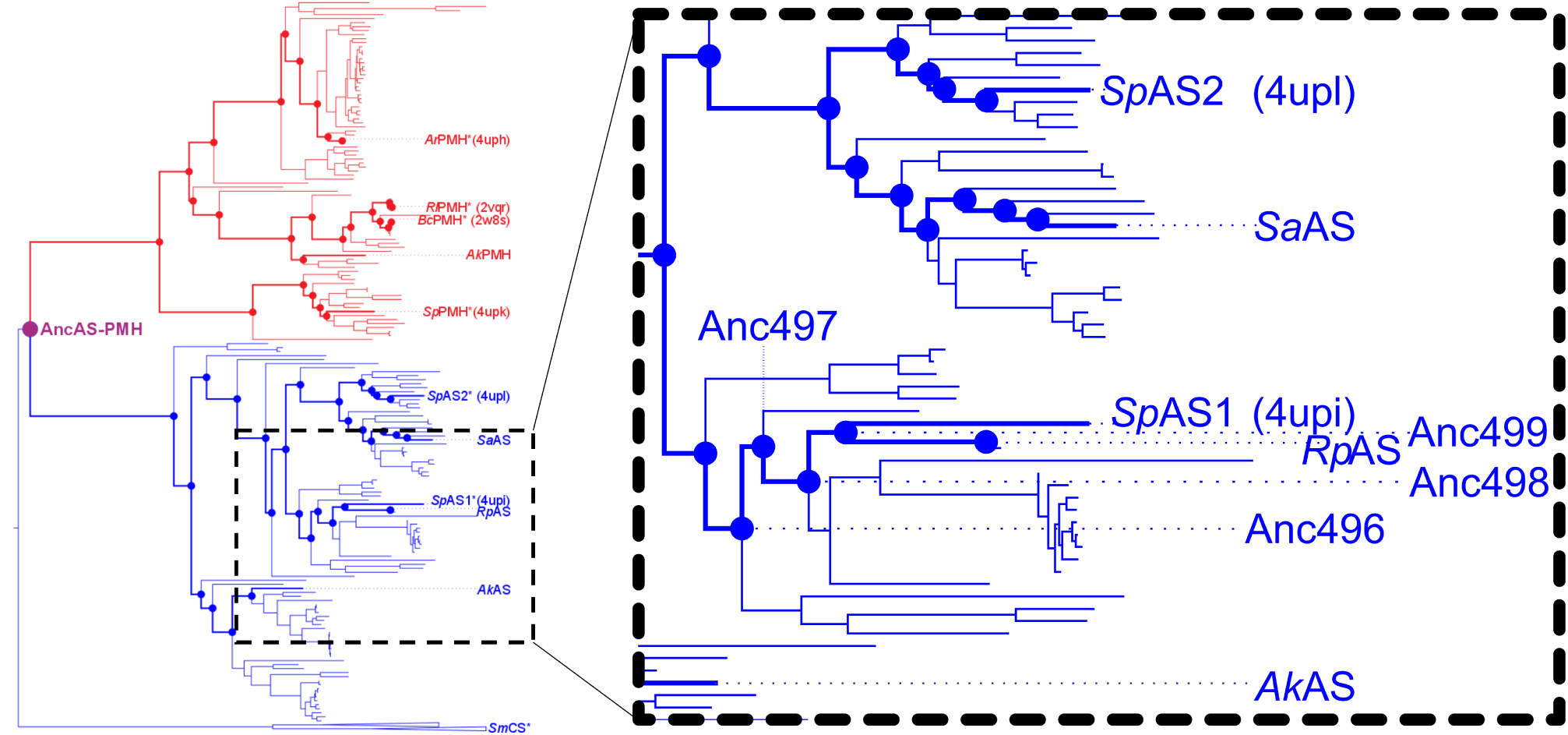
Phylogenetic tree of the AS and PMH families of the AP-superfamily leading back to the common ancestor of AS and PMH. Dots represent inferred ancestral states of the PMHs (red) and ASs (blue). Wildtype enzymes used in the study are labeled. Inferred ancestral enzymes reconstructed and used in the study are labeled in blue.

To compare the evolvability of extant and ancestral enzymes we selected three ancestral members (Anc497, Anc498 and Anc499, descending from to the presumed common ancestor of ASPMH) and four extant members (*Sp*AS2, *Sa*AS, *Ak*AS and *RP*AS) of the AS family of the previously described AP superfamily [13, 21, 60].These extant enzymes were selected to obtain a balanced representation of the AS family, and have previously been successfully expressed and purified [58] (Figure 4). Each represent a sub-group of the ASs that descend from ASPMH. We reconstructed the ancestors from *Sp*AS1 towards the root of our set of Ass until the common ancestor of ASs (see Figure 4). We selected the ancestral enzymfes to provide a range of the evolutionary history of ASs.

We successfully expressed each enzyme on the surface of *E. coli*. We confirmed expression and activity of the enzymes by observing the successful turnover of 4-NPS by intact *E. coli* cells surface displaying AS and purification of membrane fractions (Figure S1). We confirmed turnover of 4-NPS for all ancestral (Anc496, Anc497, Anc498 and Anc499) and extant (*Sp*AS2, *Sa*AS, *Rp*AS and *Ak*AS) ASs in the surface display assay. Observed enzyme activities were roughly proportional to those observed in cytosolically expressed and purified proteins.

### Creation and screening of mutant libraries

In order to investigate the comparative evolvability of ancestral and extant enzymes, we created error-prone mutant libraries of each ancestral and extant variant. We created mutant libraries through error-prone PCR using either 8-oxo dGTP or dPTP as mutagenic nucleotides to create mutant libraries with distinct mutational profiles (see Methods: Library generation methods). We aimed for a library size of 10^5^ variants to obtain a good coverage of single mutations. We tested each colony by transforming the ligation mixture and counting colonies (Table S1) and the successful insertion of each of the genes was determined by colony PCR (Figure S2).

After verifying the size and integrity of each library we transformed the remainder of the ligation mix into *E. coli* and divided the cells equally on agar plates. Sulfatase activity was monitored using the FADS with fluorescein disulphate (FDS) as a fluorescent model substrate, in microtiter plates using both FDS and 4-nitrophenol sulfate (4NPS) as substrates and on agar plates using x-sulfate (5-Bromo-4-chloro-3-indoxyl sulfate, as substrate (Figure 1).

Ancestral AS sequences were previously inferred towards the root of our set of Ass until the last common ancestor of ASs and PMHs (see Figure 4) ancestor of AS and PMH. Detailed phylogenetic analysis was performed on extant AS family members [23]. In this study we describe a parallel directed evolution campaign on three ancestral and four extant AS members for improved catalytic efficiency towards FDS. Autodisplay in *E. coli*, combined with microfluidic FADS, is used to screen and sort living *E. coli* cells presenting unique enzyme variants, with a focus on the shape of the activity distributions of enzyme variants (local fitness landscapes) obtained from FADS sorting (Figure 1).

All mutant libraries reached the target size of 10^5^ unique variants (Table S1). We determined the amino acid substitution rates for dPTP and 8-oxo dGTP libraries using Sanger sequencing. In total, we screened 14 libraries, representing two different mutational conditions: i) dPTP nucleotides (2.3±1.7 mutations) and ii) 8-oxo dGTP nucleotides (3.7±2.6 mutations) and 7 different enzymes, each either inferred ancestral or extant members of AS gr of the AP-superfamily. We verified the occupancy of the droplets by observing a sample of droplets using confocal microscopy (Figure S3). For each library we screened 10^6^ occupied droplets (λ = 0.35). As a control and to form a baseline of wild-type activity we screened 10^4^ occupied droplets of each respective wild-type enzyme in the same session using the same device for each set.

With the help of FADS up to 10^6^ variants could be tested for each library, guaranteeing a large coverage of all possible single mutations and full coverage of the targeted 10^5^ sized mutant libraries. Usage of the *E.coli* autodisplay system [25, 46] allowed for the direct recovery of recovered intact cells after screening.

All enzymes that had been previously shown to express solubly and active in cytosolic form [13, 58] also expressed well as autodisplay construct. By using FADS with living cells, it became possible to directly recovering the screened cells and plating them on agar instead of recovering and re-transforming the plasmid.

While the rates of improvement varied widely (Figure 5), the overall trend being observed showed a larger increase in improvement in sulfatases having a lower initial activity before mutation. A qualitative effect of mutation rate on enzyme evolvability can also be observed between dPTP (lower mutation rate) and 8-oxo dGTP (higher mutation rate). Here, we observed that a higher mutation rate leads to a decreased improvement in some libraries.

**Figure 5:**
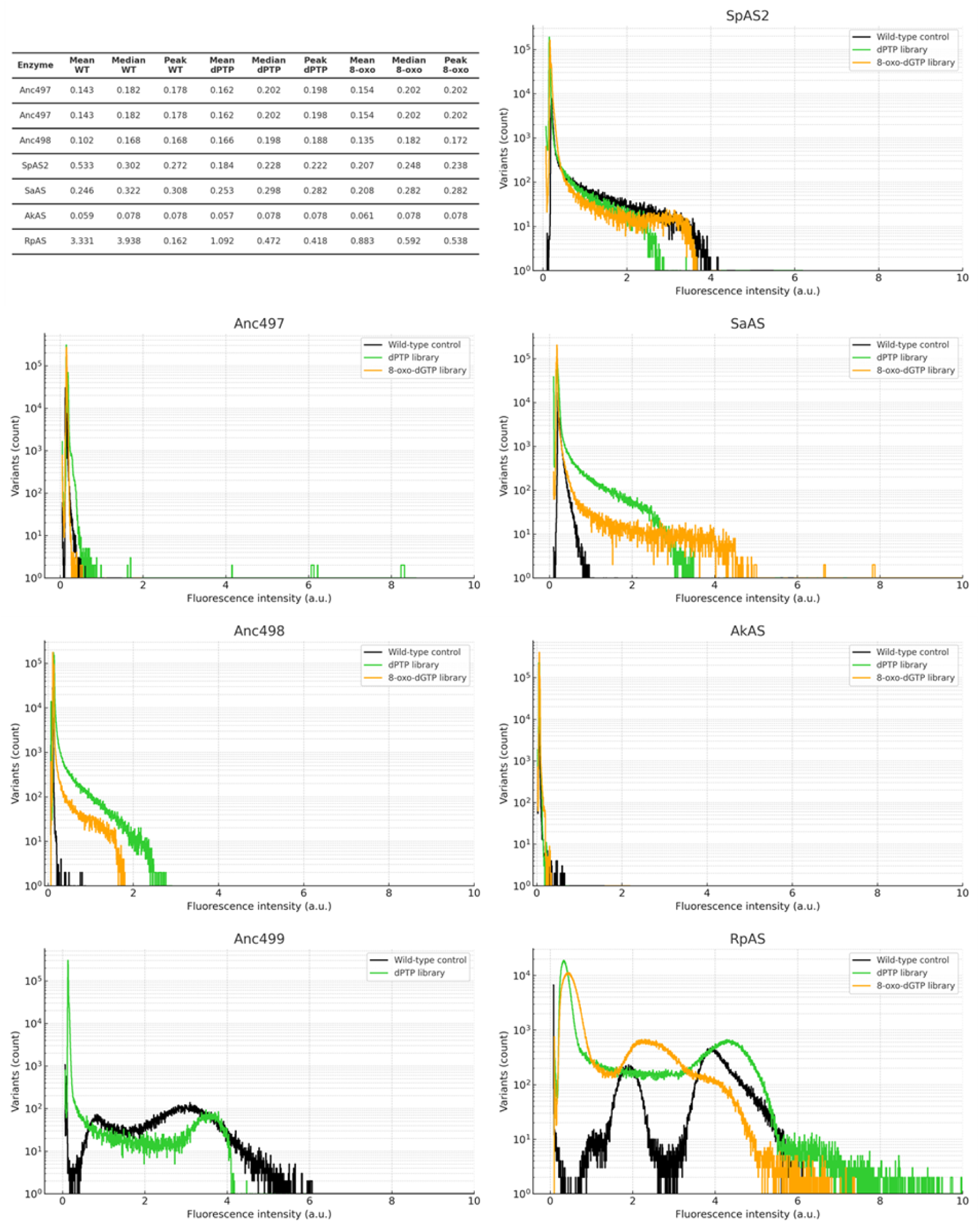
Activity distribution of each ancestral and extant enzyme. 1.000.000 individual droplets were screened using FADS. Libraries shown are dPTP (green) and 8-oxo-dGTP (orange). To measure the baseline activity of each enzyme 100.000 wild-type variants of each variant were screened (black).

The distribution of extant *SaAS* broadly followed the theorized pattern i.e. that an increased mutation rate leads to a lower number of improved variants. However, the distribution shows variants with higher activity than in the lower mutation rate library (Figure 5). This implies that variants with more than one mutation are leading to a maximum improvement, and that the coverage of variants with multiple mutations is sufficient to sort these improved variants. While the library size of 10^5^ covers each single mutant, the library does not exhaustively cover double mutations. In Anc497 and Anc498 a different trend is found. In the case of Anc497 most improved variants are seen in the dPTP library while for Anc498 most improved variants are seen in the 8-oxo dGTP library (Figure 5). This distribution sheds light on the shape of the local fitness landscape, as different mutational rates lead to a quantitatively different outcome. Remarkably for *RP*AS, the enzyme with the greatest wild-type catalytic efficiency, only minor improvements were observed in the libraries, with the majority of the pool of variants performing worse than their wild-type, in both the 8-oxo-dGTP and dPTP libraries. In the *Sp*AS2 library we observed a similar trend, with the bulk of the variants at or below wild-type level, with an extremely low proportion of improved variants. For *Ak*AS this was true for the dPTP library, although the 8-oxo-dGTP library showed a wider distribution of improved variants.

### Blue-White and agar plate screening

Blue-White screening of the cells recovered from droplets resulted in an enrichment of blue colonies (Table S3). The Blue-White screening step ensured that highly active variants were identified and selected for microtitre plate screening in the next steps.

However, Blue-White screening only provides a qualitative result and does not give any quantitative information. Therefore, Blue-White screening functions as a pre-screening method for microtitre plate screening, to ensure that all active variants are selected. Since the wild-type activity levels of some ASs were relatively low, and thus variants may appear as white due to lowered activity towards X-sulfate, even if the variant has improved activity toward FDS or 4NPS. As a result, Blue-White screening identified positives but not negatives.

Enrichment varied widely between variants. A trend was observed that dPTP libraries (low mutation rate) consistently showed a higher degree of enrichment than their 8-oxo-dGTP library counterparts (high mutation rate, see Table 1). *Ak*AS was the exception, with a higher enrichment in the 8-oxo-dGTP library, consistent with the relatively poor performance of the *Ak*AS dPTP library in screening (Figure 5). Furthermore, enrichment was higher in variants with higher initial activity, while several wild types had low activity towards X-sulfate near the detection limit. Hence, the lower enrichment may be caused by catalytic efficiency toward the X-sulfate model substrate not increasing proportionally with the catalytic efficiency towards the FDS substrate, that was used in microfluidic sorting. Thus, white colonies can still contain variants with improved catalytic efficiency toward FDS and 4NPS.

**Table 1:** Top improvements towards FDS and 4NPS compared to wild type after rescreening in 96-wells plate, including in which library the variant was found.

|  | **FDS** | **4NPS** | library |
| --- | --- | --- | --- |
| Anc497 | 4.47 | 4.83 | dPTP |
| Anc498 | 1.24 | 4.00 | dPTP |
| Anc499 | 0.74 | 1.68 | dPTP |
| *SpAS2* | 52.46 | 2.07 | 8-oxo-dGTP |
| *SaAS* | 289.17 | 5.05 | dPTP |
| *AkAS* | 58.04 | 20.59 | 8-oxo-dGTP |
| *RPAS* | 4.70 | 1.32 | 8-oxo-dGTP |

Subsequently, preliminary microtitre plate screening was carried out on variants picked from the sorted libraries. 176 picked clones of each library were screened. Variants that showed improved catalytic efficiency towards 4NPS where found in each library, while variants that showed improved catalytic efficiency towards FDS were found in each library except Anc499 (Table 1). The largest improvements were found in variants with low initial activity. When comparing ancestral enzymes to extant enzymes the most notable difference was that while all extant enzymes showed their greatest improvement towards the FDS substrate also used in FADS screening, ancestral enzymes showed their greatest improvements in catalytic efficiency over their wild-types towards the chromophoric substrate 4NPS. This difference between substrates is most pronounced in ancestral enzymes Anc498 and Anc499, and reversely extant enzymes *Sp*AS2 and *Sa*AS. The biggest improvements towards both 4NPS and FDS were observed in ancestral enzymes Anc497 and Anc498 for both FDS and 4NPS, and extant *Sa*AS for activity towards FDS and *Ak*AS for activity towards 4NPS, respectively. Coincidentally, these four enzymes showed low initial activity towards FDS. Although only anecdotal, these results hint that variants with lower initial catalytic efficiency may ’catch up’ and exceed wild-type enzymes with high initial catalytic efficiency even after only a few rounds of directed evolution.

### Evolvability of ancestral enzymes

Ancestral enzymes, with their enhanced thermostability and substrate promiscuity, are suggested as promising candidates for directed evolution [61, 26, 38]. With the growing accessibility of synthetic DNA, experimental exploration of ancestral enzyme properties has become feasible. To enable direct parallel comparison of ancestral and extant proteins, the mutant libraries were concurrently created using identical methods and backbones.

We assessed the evolvability traits of four extant and three ancestral ASs belonging to the AS-PMH branch of the alkaline phosphatase superfamily (Figure 4). Our aim was to correlate the evolutionary status of the enzyme with its evolvability and robustness. Leveraging FADS, we conducted high-throughput parallel screening of both modern and ancestral enzyme libraries, as well as preliminary rescreening of recovered variants.

The variation in mutation rates highlights the distinct shape of the local fitness landscape for each enzyme. Earlier studies on malate dehydrogenase and lactate dehydrogenase enzymes already showed that a single crucial mutation can switch substrate specificity and greatly impact enzyme activity [33]. Given that most mutations are either neutral or destabilizing, stabilizing mutations may be necessary to achieve a functional protein with improved or distinct functions. This contrast may explain the disparity between enzyme backbones capable of attaining novel functionality in one step and those needing additional backbone stabilization to achieve novel function [62, 37].

As expected, the observed mutational landscape varied depending on the enzyme and mutational regime. Naturally, increasing the mutation rate would flatten the curve and possibly widen the activity distribution. Given that most mutations are either neutral or detrimental, accumulating multiple mutations increases the likelihood of encountering a deleterious mutation that impairs variant function.

In addition to an increase in impaired variants, one would expect an increase highly improved variants if multiple point mutations are needed to achieve a substantial increase in function. These mutations can either serve as compensatory mutations, necessary to counteract stability loss or folding issues, or as additional mutations directly facilitating changes in function. By introducing multiple mutations in a single round, variants requiring compensatory mutations or epistatic interactions become available. However, each mutation may also cause the enzyme to be non-functional. Here a balance between the possibility of jumping evolutionary ’ratchets’ and avoiding the accumulation of deleterious mutations appears. These mutations can be either compensatory mutations, which are required to offset thermostability loss or folding issues or additional mutations. Additional mutations are required to directly facilitate changes in function. Often, while enzymes with different functions differ in many residues, only a few mutations are responsible and sufficient for a change in function [33, 63]. When comparing this phenomena, in which a few mutations are able to greatly shift the activity of an enzyme, to our library screening we observed a similar phenomenon especially in the case of *Sa*AS and *Ak*AS (Figure 5). An interesting case is observed in Anc497, where the dPTP library led to greatly improved variants up to five-fold. Yet, the 8-oxo-dGTP library led to much less improved variants, with the top variant showing only half the improvement of most active dPTP variants, as well as showing significantly fewer improved variants (Figure 5). These results are consistent with a specific fitness landscape for each variant, such that the optimal number of mutations to achieve an improvement may vary between enzymes. The histograms indicate that ancestral libraries yielded on average fewer improved variants than the extant enzyme *Sa*AS (Figure 5). However, the ancestors yielded more improved variants than *RP*AS, *Sp*AS2 or *Ak*AS. Interestingly, in most ancestral enzymes the lower mutation rate dPTP library yielded more improved variants, while the higher mutation rate 8-oxo dGTP library yielded more improved variants in extant enzymes.

Furthermore, when improved variants were screened in microtiter plate format, increases in catalytic efficiency towards both FDS **3a** as well as 4NPS **1a** (that was not used for microfluidic screening) were found. This effect was most pronounced in ancestral variants Anc497, Anc498 and Anc499, in which the increase cataltyic effiency towards 4NPS was more substantial than higher than the increase in catalytic efficiency towards FDS.

Overall, these results show that ancestral enzymes can show promise as starting points for directed evolution. When it came to extant enzymes, our results showed that the enzymes with greatest starting kinetic parameters showed the least overall improvement, with *Sa*AS outperforming these initially more fit enzymes. Notably, libraries with a lower mutation rate showed a greater proportion of highly improved variants in ancestral libraries, whereas higher mutation rates showed greater proportion of highly improved variants in extant libraries. This characteristic makes ancestral enzymes especially interesting when screening is limited to lower throughputs due to limitations such as substrate or product detection.

## Conclusion

In this study, we integrate microdroplet sorting, autodisplay technology, and Ancestral Sequence Reconstruction (ASR) on the alkaline phosphatase superfamily, focusing on the Aryl sulfatases (AS) and phosphonate monoester hydrolases (PMH) [13]. Our objectives were (i) to systematically investigate the evolvability and robustness of ancestral proteins, (ii) to establish a correlation between catalytic efficiency enhancements observed through high-throughput methods and those observed in cytosolically expressed, purified proteins, and (iii) to gain insights into the differences and similarities in the local fitness landscapes of various extant and ancestral As variants.

We demonstrated that cell-surface-displayed enzyme libraries, grown within droplets, are effective for sorting large libraries—each containing over 10^5^ variants—in parallel within a practical time-frame, as well as applying droplet microfluidics to screen proteins derived from ancestral sequence reconstruction for the first time.

We conclude that, in addition to screening of directed evolution libraries, microfluidic FADS sorting and recovery of autodisplayed proteins can be utilized for a variety of applications in high-throughput screening and sorting of proteins. An example would be the screening of synthetic libraries of proteins derived from multiplexed gene synthesis [64, 65].

When comparing the variant activity histograms of the tested Arylsulfatases (ASs), we found that evolvability was maximized under a lower mutation rate for ancestral enzymes while the extant enzymes benefited from a higher mutation rate. On the other hand, for the ancestral libraries the lower mutation rate dPTP library resulted in a greater proportion of highly improved variants (Figure 5), demonstrating a qualitative difference in the ideal mutation rate of different enzymes.

In the future, this study could be followed up with a much more thorough biochemical characterization of improved variants, including detailed kinetics of purified enzymes and elucidation of crystal structures, as well as introduction of the mutations found in the screening into corresponding positions in different enzyme backbones.

Reconstructed ASs showed a propensity towards needing fewer mutations for improved function. Therefore, ancestral enzymes, particularly in the context of ASs, can serve as valuable initial scaffolds for rapidly achieving significant enhancements in enzyme activity.

Additionally, "imperfect reconstruction"—stemming from less than 100% certainty in assigning residues at ambiguous sites [66]—can be refined through high-throughput directed evolution. Comprehensive coverage of single mutations includes all one-aminoacid-off alternative reconstructions, allowing for the selection of variants that enhance functionality.

An initial directed evolution campaign with multiple (both ancestral and extent) enzymes increases the odds of finding success, as the candidate with optimal initial catalytic efficiency may not have traversable, single/double mutation route towards desired function.

Accuracy versus throughput is a consideration throughout the experiment. This leads to a funnel-like progress where high-throughput screening is used to select improved variants from a large pool, which are then further characterized using lower throughput microtitre plate screening of variants and finally purified protein.

As ASs typically show promiscuity towards other substrate classes the concept of a low initial activity construct with a broad fitness landscape can be extended towards different enzymes with a promiscuous activity towards the desired reaction.

It should be cautioned screening towards a limited number substrates only investigates *biological* specificity and not *intrinsic* specificity [40]. As of such, while ancestral enzymes may be more *biologically* generalist, a trend linking between ancestrally reconstructed sequences and reduced *intrinsic* specificity has not been shown, and might not be expected [40].

These results also open up several lines of inquiry for future work. A first line of inquiry is a longer directed evolution campaign starting from improved variants of the best performing ancestral and extant enzyme. Another avenue is expanding the screening from sulfate esters (4NPS and FDS) towards substrates for promiscuous activity towards phosphonate esters and phosphate mono- and di-esters. A further path is screening in parallel through a larger array of conditions such as higher mutation rates or differently biased libraries.

## Author contributions

The experimental design was largely performed by B.D.G.E., T.S.K., E.B., J.J., F.H., and B.V. B.D.G.E, M.H., J.M.H., A.L., C.D., T.S.K. and B.V. performed experiments. B.D.G.E, T.S.K., A.L. and E.B drafted the first major version of the manuscript with improvements and directions from all other authors.

## Data availability

Data for this article, including FADS screening data and the DNA sequences of the proteins used are available at Zenodo at https://doi.org/10.5281/zenodo.16812894

## Acknowledgments

E.B.B. and A.L. received funding from Volkswagen foundation grant code 98183. E.B.B. from HFSP (Human Frontiers of Science Programme, RGP0006/2013 and RGP004/2023); DFG (Deutsche Forschungsgemeinschaft BO-2544/20-1;503272152. B.E. received funding from the EU under the Horizon 2020 Research and Innovation Framework Programme No. 722610. TSK was supported as an EU H2020 Marie Skłodowska-Curie Fellow No 750772. FH is an ERC Advanced Investigator (695669).

