## Supplementary Information for "Studying the evolutionary potential of ancestral aryl sulfatases in the alkaline phosphatase family with droplet microfluidics"

### Supporting Information

Outer membrane fraction protein preperation: 80 mL of LB-medium was inoculated with each *E. coli* with a pBAT-AT vector containing each enzyme variant. Cultures were incubated while shaking until OD was approximately 0.8 and induced with 0.1% arabinose. After overnight incubation at 25 °C samples were split into two 40 mL portions. All samples were centrifuged for 10 min at 3850 xg and supernatant was discarded. Samples treated with proteinase K (+) were resuspended in 1 mL PBS, 12.5 µL proteinase K (5mg/mL) and were incubated at 37 °C for 1 hour, the reaction was then stopped by adding 5 mL Tris/HCl + 10% FCS. The solution was centrifuged for 5 min at 3850 xg and supernatant was discarded. Next both the proteinase K treated samples (+) and untreated samples (-) were resuspended into 1,5 mL Tris/HCl solution 0,2 M at pH 8.0. 0.1 mL sucrose 1M, 0.1 mL EDTA 10 mM, 0.1 mL lysozyme (10mg/mL) and 3.2 mL ultra-pure water. The solution was incubated 10 min at room temperature. Subsequently 50 µL PMSF (100 mM, in Isopropanol), 10 µL Aprotinin (10mg/mL, in HEPES-buffer), 5 mL extraction buffer (2% Triton X100, 50mM Tris/HCl, 10 mM MgCl2), and 100 µL DNAse (1mg/mL) was added, and the mixture stored for 30 m on ice. The solution was then centrifuged at 3850 xg and the pellet was discarded and the solution transferred to a centrifuge tube. THe supernatant was centrifuged for 10 min at 18000 rpm, Sorvall®. The supernatant was discarded, the pellet was resuspended in 10 mL millipore water and centrifuged for 10 min at 18000 rpm. The supernatant was discarded, the pellet resuspended in 1.5 mL Millipore water and tranferred into a 1.5 mL reaction tube. This solution was centrifuged for 10 min at 15000 xg, the supernatant discarded, and the pellet resuspended into 20-100 µL ultra-pure water proportional to pellet size. The membrane fractions were then analyzed using SDS-page.

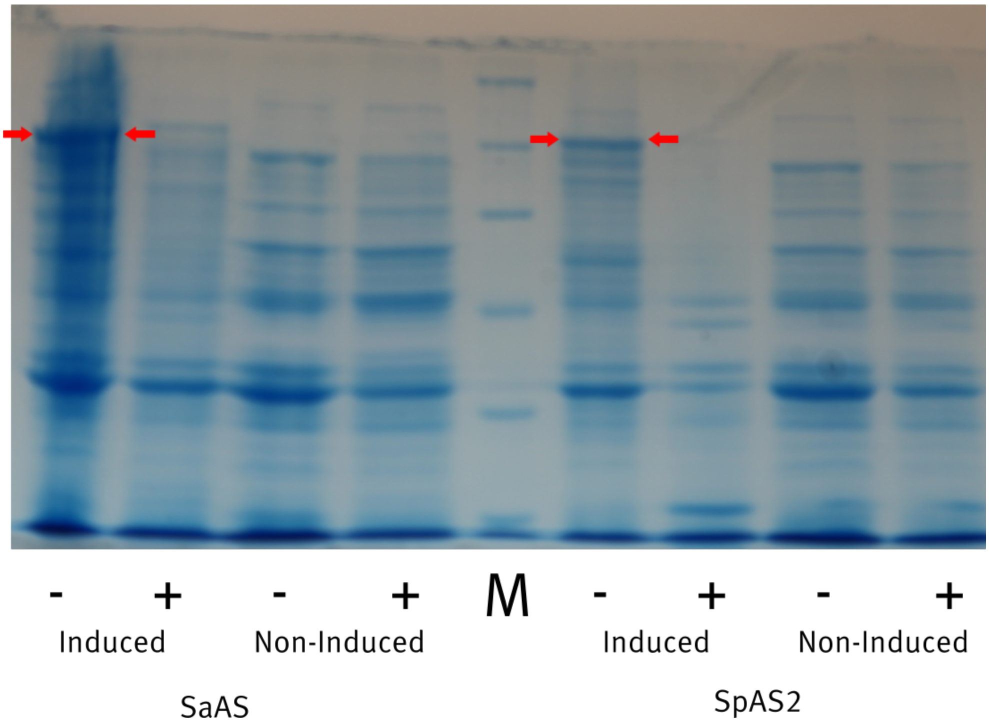


Figure S1: Purified membrane fractions of enzymes *Sa*AS and *Sp*AS. Showing both induced and non-induced cells with (+) and without (-) proteinase K treatment.

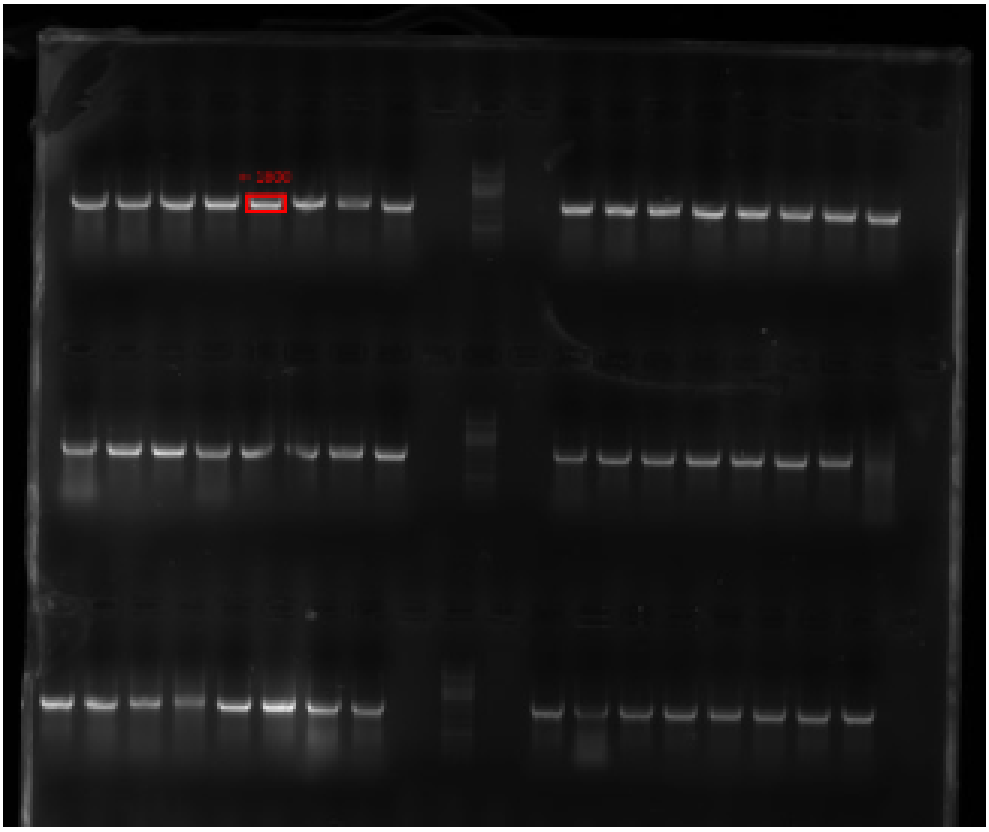

Figure S2: Colony PCR showing the successful insertion of fragments in 8 colonies tested from each library.

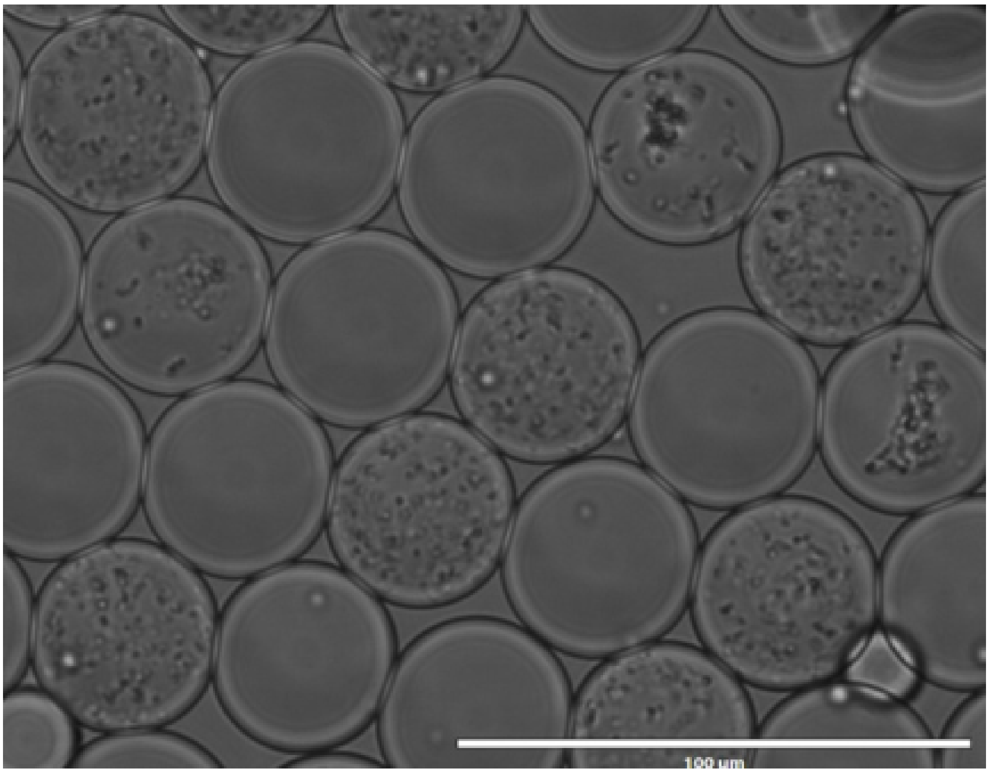


Figure S3: Confocal microscope image showing 40 pL microdroplets with and without cell culture after encapsulation of single cells and overnight incubation.

| **Library** | **estimated size** |
|---|---|
| Anc497 8-oxo dGTP | 228.000 |
| Anc497 dPTP | 426.000 |
| Anc498 8-oxo dGTP | 462.000 |
| Anc498 dPTP | 396.000 |
| Anc499 8-oxo dGTP | 270.000 |
| Anc499 dPTP | 450.000 |
| *Sp*AS2 8-oxo dGTP | 270.000 |
| *Sp*AS2 dPTP | 1.926.000 |
| *Sa*AS 8-oxo dGTP | 750.000 |
| *Sa*AS dPTP | 402.000 |
| *Ak*AS 8-oxo dGTP | 360.000 |
| *Ak*AS dPTP | 1.236.000 |
| *RP*AS 8-oxo dGTP | 822.000 |
| *RP*AS dPTP | 1.218.000 |

Table S1: Estimated sizes of each generated library, library sizes were determined by extrapolating from dilution plates.

| | **"Age"** | **Tm** |
|---|---|---|
| Anc497 | 0.72 | 57.4 |
| Anc498 | 0.75 | 49.6 |
| Anc499 | 0.78 | 48.6 |
| *Sp*AS2 | 0.98 | 51.7 |
| *Sa*AS | 1.00 | 49.5 |
| *Ak*AS | 0.68 | 45.6 |
| *RP*AS | 0.90 | 47.05 |

Table S2: Thermostability of extant and ancestral wild-types.

| | **oxo** | | | **ptp** | | |
|---|---|---|---|---|---|---|
| | **Blue** | **White** | **%** | **Blue** | **White** | **%** |
| Anc497 | 3 | 147 | 0.020 | 6 | 331 | 0.018 |
| Anc498 | 45 | 2500 | 0.018 | 96 | 2500 | 0.037 |
| Anc499 | 439 | 2000 | 0.180 | 722 | 2000 | 0.265 |
| *Sp*AS2 | 38 | 2500 | 0.015 | 1239 | 2500 | 0.330 |
| *Sa*AS | 32 | 170 | 0.158 | 89 | 223 | 0.285 |
| *Ak*AS | 43 | 308 | 0.123 | 12 | 273 | 0.042 |
| *RP*AS | 858 | 2800 | 0.235 | 2084 | 1500 | 0.581 |

Table S3: Percentage of enriched colonies after Blue-White screening.
